# Isolation of Cyclotide Gene from Solanaceae Family and Its Bioinformatics Analysis

**DOI:** 10.1101/2025.06.02.657358

**Authors:** Muhammad Ahmad, Mehnaz Ghulam Hussain

## Abstract

Cyclotides are highly tolerant of sequence variability, aside from the conserved residues forming the cystine knot. Cyclotides have attractive natural movement capabilities including cytotoxicity, harmfulness to HIV, antimicrobials, insecticides, uterine contraction and inhibition of proteases, making them very important in pharmacy and horticulture. The medicinal cyclotides may be able to reimburse the safety of antimicrobials, which is a noticeable problem worldwide today. The present study was focused on the isolation of genes that encoding cyclotide, from plant species. DNA was isolated by CTAB method. The gene was amplified by PCR and after agarose gel electrophoresis confirmation that gene was sequenced followed by bioinformatics analysis. Petunia showed 90% to 97% identity with cds of reference sequence and 27 amino acids were identical in multiple alignment of protein sequence. In addition to providing information for the future investigation of Solanaceae species for cyclotides, this work will aid in understanding the plant families that produce cyclotides.

## Introduction

Plant-derived cyclic peptides known as cyclotides have three conserved disulfide bridges that form a stable cystine knot motif and a distinctive head-to-tail circular backbone. Cyclotides are desirable candidates for therapeutic treatments due to their structure, which also confers resistance against enzymatic, chemical, and thermal destruction (1). Further, their architecture can function as a molecular scaffold to produce peptide-derived medications with enhanced stability and bioactivity because of their tiny, stable structure.

Cyclotides show a wide range of pharmaceutical action including anti-microbial, anti-HIV, cytotoxic, hemolytic, and anti-inflammatory activities (2). These multi-functional peptides are a part of many plant defense mechanisms that can protect plants from pathogens and environmental pressures (3). Although cyclotides have been studied extensively in families such as Violaceae, Rubiaceae, and Cucurbitaceous, it remains to be seen how many these unique sequences exist in the Solanaceae family of plants, which contains over 3,000-4,000 species (4). Solanaceae contains some economically important plants such as Petunia and Solanum, which may represent an interesting source of novel cyclotide variants with important biotechnology (5).

Recent strides in bioinformatics, transcriptomics and high-throughput sequencing have enabled the discovery of cyclotide-like genes in non-traditional plant families, such as Poaceae and Fabaceae (6). However, while many studies involving cyclotides have examined *Solanaceae* species from other standpoints, the systematic isolation and characterization of cyclotides from this family have only recently commenced, despite their immense potential for drug design and agricultural purposes (7). Cyclotides, such as Kalata B1 from Viola odorata have been shown to elicit therapeutic responses in infection treatments, inflammation, and cancer (8) and the process by which cyclotides are produced is thought to involve cyclization via asparaginyl endopeptidase (AEP) that is still being investigated for optimal recombinant production (9).

Moreover, bioinformatics tools help to make predictions regarding cyclotide genes, to analyze the genes in regards to their conserved domains, and to study the evolutionary relationships of families of plants. The computational approaches, like database mining and molecular modeling have identified cyclotide-like sequences in maize and other crops, indicating a broader phylogenetic distribution (10). However, the biosynthetic pathways, the expression patterns, and the biological functions of cyclotides derived from Solanaceae are largely unknown.

This research aimed to Isolate and characterize the cyclotide genes from the Solanaceae family, conduct bioinformatics analyses to predict structural and functional properties, examine their phylogenetic relationships with known cyclotides. This investigation will help discover new cyclotides and their potential uses in medicine and biotechnology by using experimental and computational method.

## Materials and Methods

### Collection of selected plant

Designated (Red Petunia, Blue Petunia, White Petunia) plants have been bought from the Faisalabad nursery in Pakistan and were kept at the required temperature and environment, shielded from water and other conditions that could be the reason for spoiling, until the DNA was extracted.

### DNA Extraction and Isolation

The CTAB technique (11, 12), which calls for 70% ethanol, chloroform isoamyl alcohol, and CTAB buffer, was used to isolate the DNA. NaCl: 8.19 g, EDTA: 0.584 g, Tris Base: 1.21 g, CTAB: 2 g, β-mercaptethanol: 0.2 ml, and distilled water: 80 ml made up the CTAB buffer. To make the CTAB buffer solution, 25 milliliters of distilled water were dissolved in Tris-base, and 0.1N HCl and NaOH were used to bring the pH down to 8. A measuring cylinder was then taken and all the measured parts were poured into it and up to 80 ml of volume was generated. On a magnetic stirrer, the solution was transferred to a beaker and allowed to further dissolve. Due to its sensitivity to light, the fully prepared solution was stored at room temperature, wrapped with aluminum foil. Following that, washed leaves from particular plants were extracted and ground into a thin paste. During the grinding process, 1000 μl of CTAB buffer was added. We collected and labeled a 0.2 g sample. A dry bath was used for 30 minutes at 65 degrees Celsius after 700 μl of CTAB buffer was added. To enable the sample to mix with CTAB, the sample was vortexed every two minutes throughout incubation. The topmost layers containing DNA were pipetted out following the addition and vigorous mixing of an equivalent volume of chloroform: isoamyl alcohol (24:1). After that, the mixture was centrifuged at 13,000 rpm for 10 minutes. After 30 minutes at -20 degrees Celsius, add 0.6 volume of isopropanol, and centrifuge for 10 minutes at 13,000 rpm. After removing the pellet and discarding the supernatant, 200 μl of 70% ethanol was centrifuged for eight minutes at 13,000 rpm. Pellets were air-dried and the supernatant was thrown away once again. The pellet was then stored at -20 °C after being dissolved in 50 μl of nuclease-free water.

### Primers Designing

For the PCR amplification of cyclotide genes (13), particular Eagle F1 (Petunia Type) reverse and forward primers created by Justbio.com were required. Primers were designed using online software (http://www.justbio.com). First, the gene’s starting and stop codon sequencing sections were chosen, and the information was retrieved from the previously published sequence in GenBank (NCBI) (https://www.ncbi.nlm.nih.gov/genbank/) (14). Initially, the Cleaner hosted tool (15) was used to remove any extraneous test marks (spaces or gaps) from the sequences. Next, the parameters of primers such as molecular weight, melting temperature, GC content etc. were calculated by the OligoCalc tool (https://www.biosyn.com/Gizmo/Tools/Oligo/Oligonucleotide%20Properties%20Calculator.htm) (16). The Complementor tool (https://www.bioinformatics.org/sms/rev_comp.html) was used to acquire the DNA’s complementary reverse sequence and the Primer Aneal tool, which uses primers to hybridize on the relevant cDNA or DNA template sequence, made it simple to search and visualize. Then ten microliters of primer and ninety microliters of deionized water were placed in an Eppendorf.

**Table 1.**
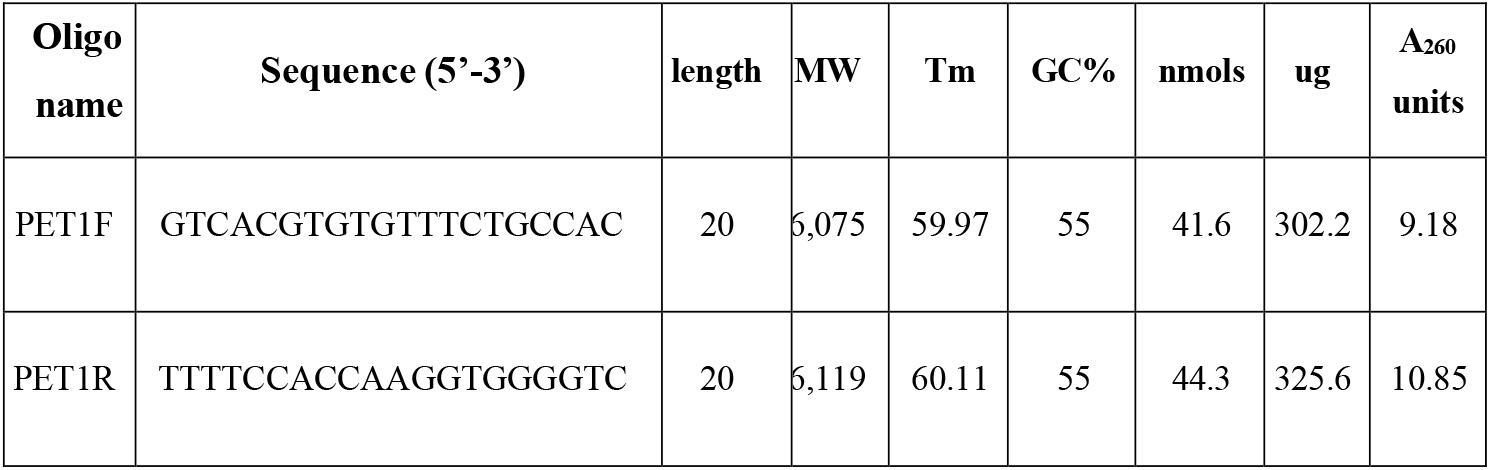
Primers Details.

### PCR Amplification

A PCR mixture was made and placed in the Thermo Cycler equipment to amplify the relevant template, the computer was instructed to conduct the PCR cycle under standard circumstances and the samples were run on a 1.2 percent agarose gel to confirm the PCR results. This was followed by column purification of DNA fragments after PCR using a kit method (Gel Purification Mini KitCat, No.: 001-1 FAGPK (200 Preps).

### Agarose Gel Preparation

An electric balance was used to weigh 0.4 g of agarose powder, which was then added to the flask having 40 cc of 1X-TAE buffer. As DNA size determines gel concentration so, the separated DNA bands were smaller at higher gel concentrations and vice versa. After that, it went microwave-heating for 1:30 minutes until it was fully dissolved and prior it being cast onto the tray, it was allowed to cool. To prepare the tray, the comb was placed within the tray, and then gel was poured inside to prevent bubbles from forming and impeding the flow of DNA molecules. To allow the gel to polymerize, it was left at room temperature for 20 to 30 minutes and as soon as it solidified, the comb was taken out. However, the tank was filled with 1X-TAE buffer to the fullest extent possible and the gel tray was inserted into the tank that was filled. After loading the sample and ladder, the device was turned on and maintained at 90 volts for 40 minutes. The gel was then taken out of the device and left in ethidium bromide for 20 minutes (17).

### Extraction and Purification

Prior to extraction from the agarose gel containing the necessary gene dry bath sequence, the temperature was set at 55 °C. Using a clean scalpel, excess agarose gel was cut off in order to digest the agarose gel that contained the relevant DNA fragments and shrink the gel slice (The ideal gel slice size was about 200 mg). Three volumes of FAGP Buffer were applied; for instance, 600 μl of FAGP Buffer was added to dissolve 200 mg of gel slice and following a 10-to 15-minute incubation period at 55° C, each tube was vortexed every five to seven minutes until the gel slice was completely dissolved (the vortex’s breakdown increases gel during incubation) and 800 μl of the sample combination was transferred to the FAGP Column. After centrifuging it for 30 seconds and discarding the flow through, the FAGP column was put back into the collection tube and 750 μl of wash buffer (ethanol) was added to the FAGP column. After discarding the flow-through, the FAGP column was put back into the collection tube and left there for 30 seconds. To dry, centrifuged the FAGP column for an additional two minutes and to elute the DNA in the elution tubes, the FAGP columns were centrifuged for one minute, after the elution buffer was periodically added to the membrane center of the column in the final varied amounts. The columns were then permitted to stand for three to four minutes and agarose gel electrophoresis was used to confirm the purity of the DNA.

### Sequencing and Bioinformatics Analysis

Sanger’s method (18) was used to sequence the required and separated genes, and samples were forwarded to CAMB Lahore. Effective bioinformatic web tools from NCBI were used to perform sequence analysis for gene characterization, phylogenetic investigations, and genetic variant comparison.

## Results

### Selected Plants

To separate the cyclotide gene from the plants, red, white, and blue petunia plants (figure 1) were easily obtained from the nearby Faisalabad nursery were chosen.

**Figure 1.**
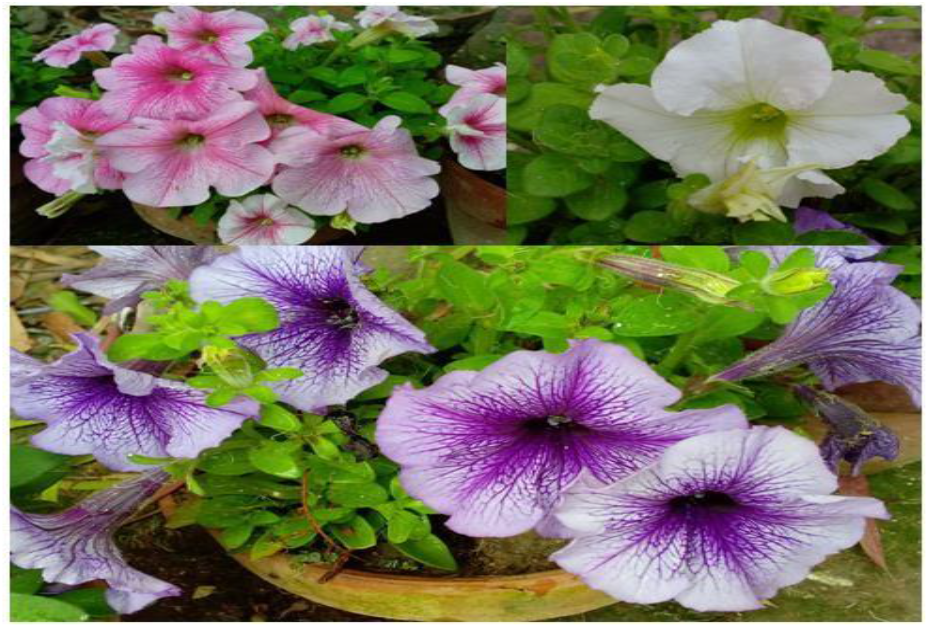
Petunia Species (Solanaceae Family)

### Isolated DNA Bands on Gel Electrophoresis

The isolated DNA of the whole sample was run on the gel one after another and the DNA isolation was effective and results of red petunia are shown in (figure 2) while is the (figure 3) is of blue and white petunia.

**Figure 2.**
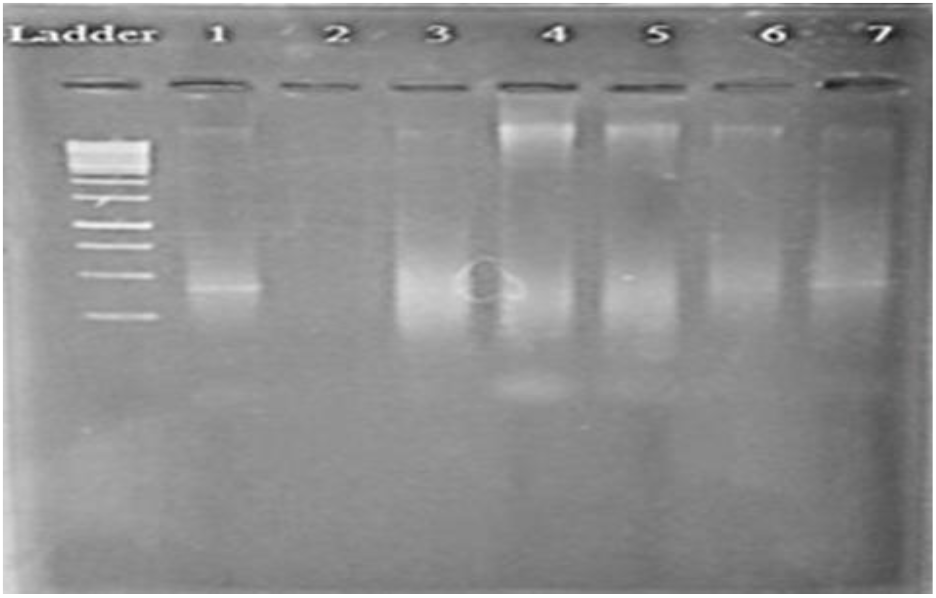
Wells 4, 5, 6 DNA band of Red *Petunia* of Solanaceae Family Plant.

**Figure 3.**
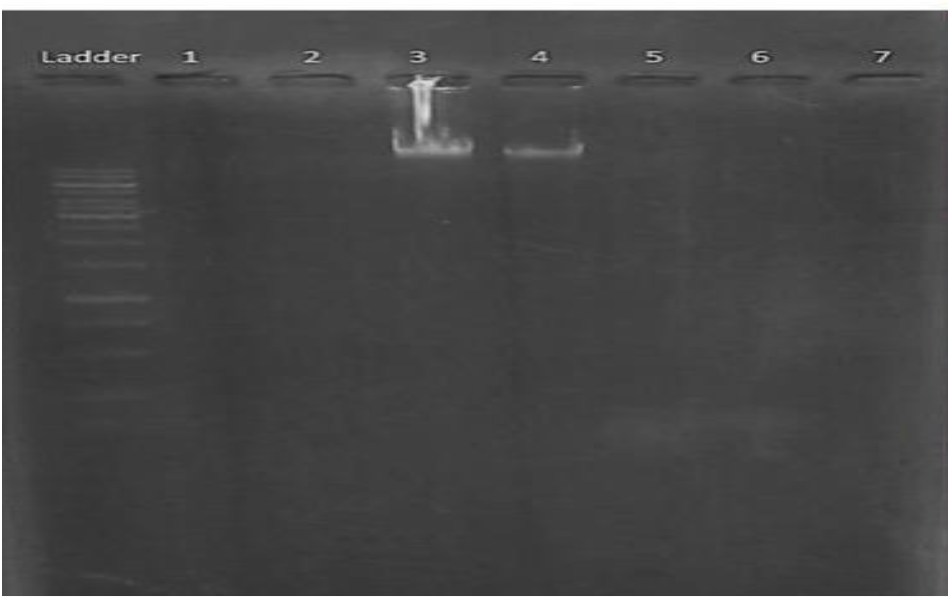
DNA bands from blue (W3) and white (W4) *Petunia* Solanaceae Family

### Primers Designing

For the amplification of AEP gene specific reverse and forward primers were designed from NCBI and online bioinformatics tools were used to design primers by using different hosted tools of this site. First, a segment of the gene was chosen whose start and stop codon regions were conserved, as confirmed by cluster alignment and GENE BLAST. This information was taken from a sequence that had already been reported from GenBank (NCBI) (figure 4). Then 10 primers were displayed by Blast, but one was chosen because its Tm and GC% looked ideal.

**Figure 4.**
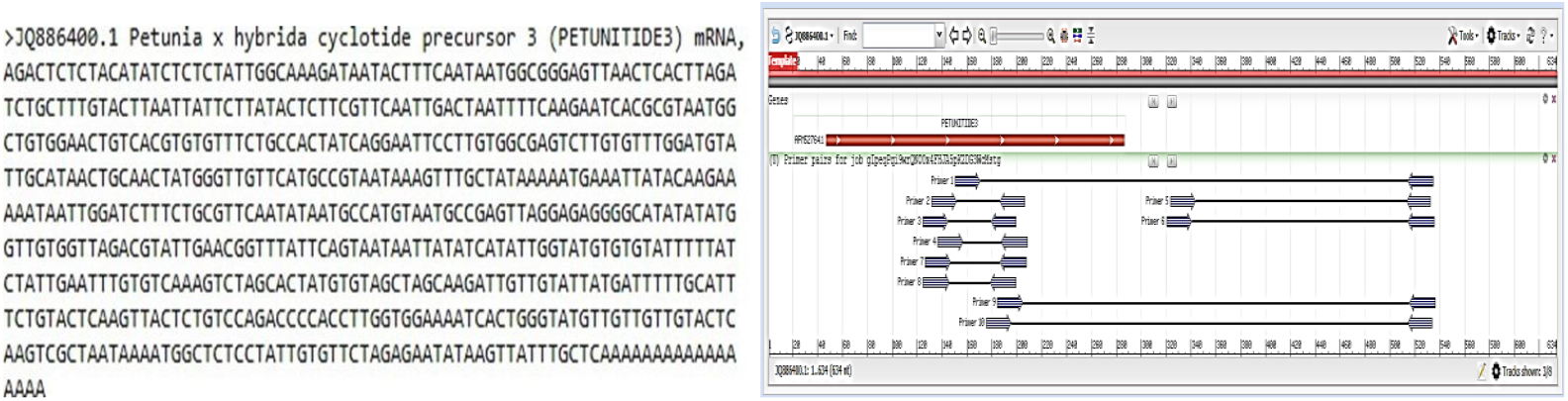
Complete CDS sequence of Petunia x hybrid cyclotide precursor 3 mRNA and graphical view of Primer

### PCR Purification and Gel Electrophoresis

A kit approach called FavoPrepTM, Gel purification Mini kit Cat. No: FAGPK 001-1 (200 Preps) was used to purify DNA fragments following PCR. Using a clean, sterilized scalpel, the gel was cut from the area containing the desired gene fragments. The gel was then placed on a UV illuminator with the appropriate UV protection and safety measures in place. The PCR-amplified gene-containing incised gel segments were selected, stored in an Eppendorf container, and the amount of gel was weighed in milligrams. Favorprep gel purification mini kit FAGPK 001-1 (200 preps) was used to purify the products (figure 5).

**Figure 5.**
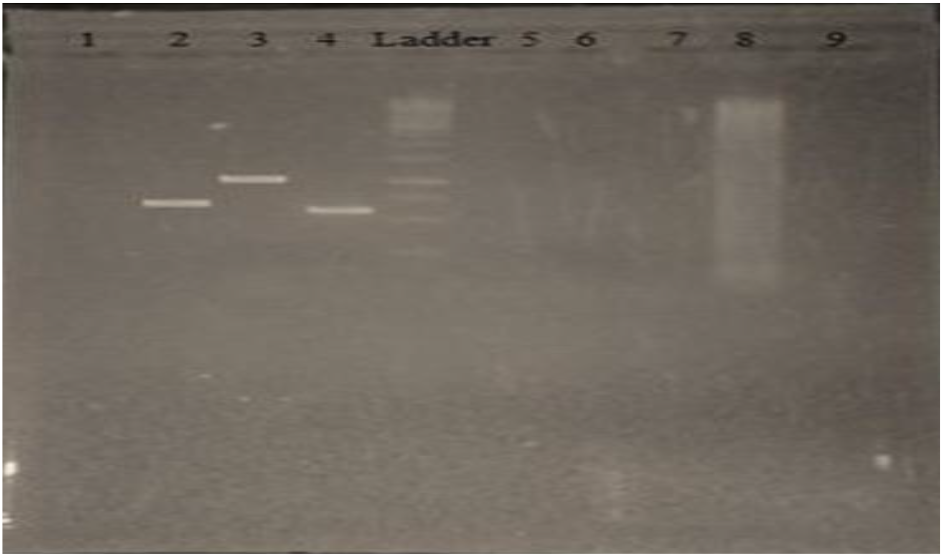
PCR Product of the Red, Blue and White *Petunia* in 2, 3, 4 wells

### Sequence Analysis and Alignment

Sequences of cyclotide genes belonging to Red, White and Blue Petunia were obtained in duplicates. After getting sequences of cyclotide gene isolated, alignment was done in next step of obtained sequences with the reference sequence by using BLAST, NCBI and other bioinformatics tools.

### Alignment of Blue Petunia Isolates with Reference Sequence (JQ886400)

First of all, alignment of isolated sequence from Blue *Petunia* with reference sequence was done and it showed 90% identity with cds of reference sequence and the alignment of isolated sequence of Blue *Petunia* with reference sequence is shown in (figure 6).

**Figure 6.**
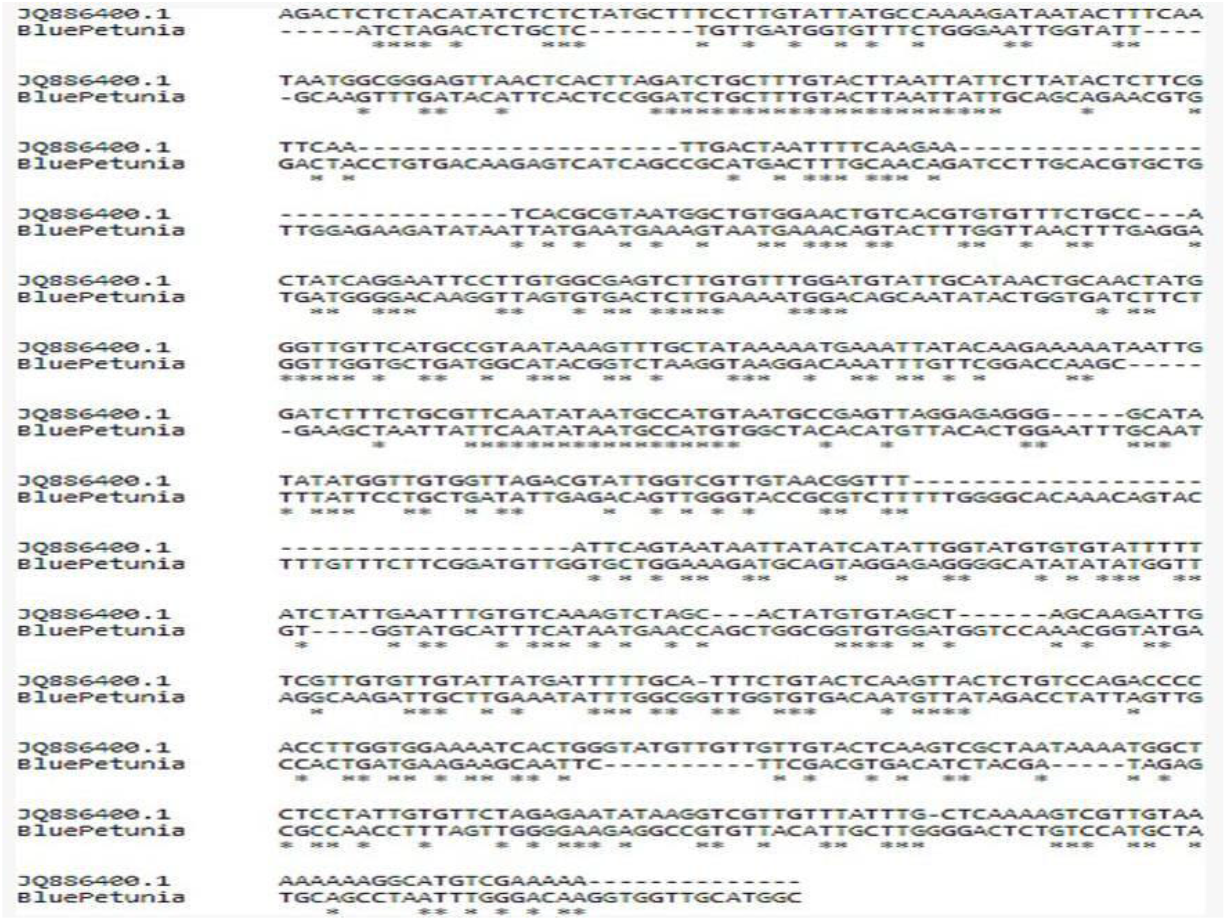
Alignment of Blue *Petunia*

### Alignment of Red Petunia Isolates with Reference Sequence (JQ886400)

Alignment of isolated sequence from Red Petunia with reference sequence was done and it showed 93% identity with cds of reference sequence. Thus, the alignment of isolated sequence of Red Petunia with reference sequence is shown in (figure 7)

**Figure 7.**
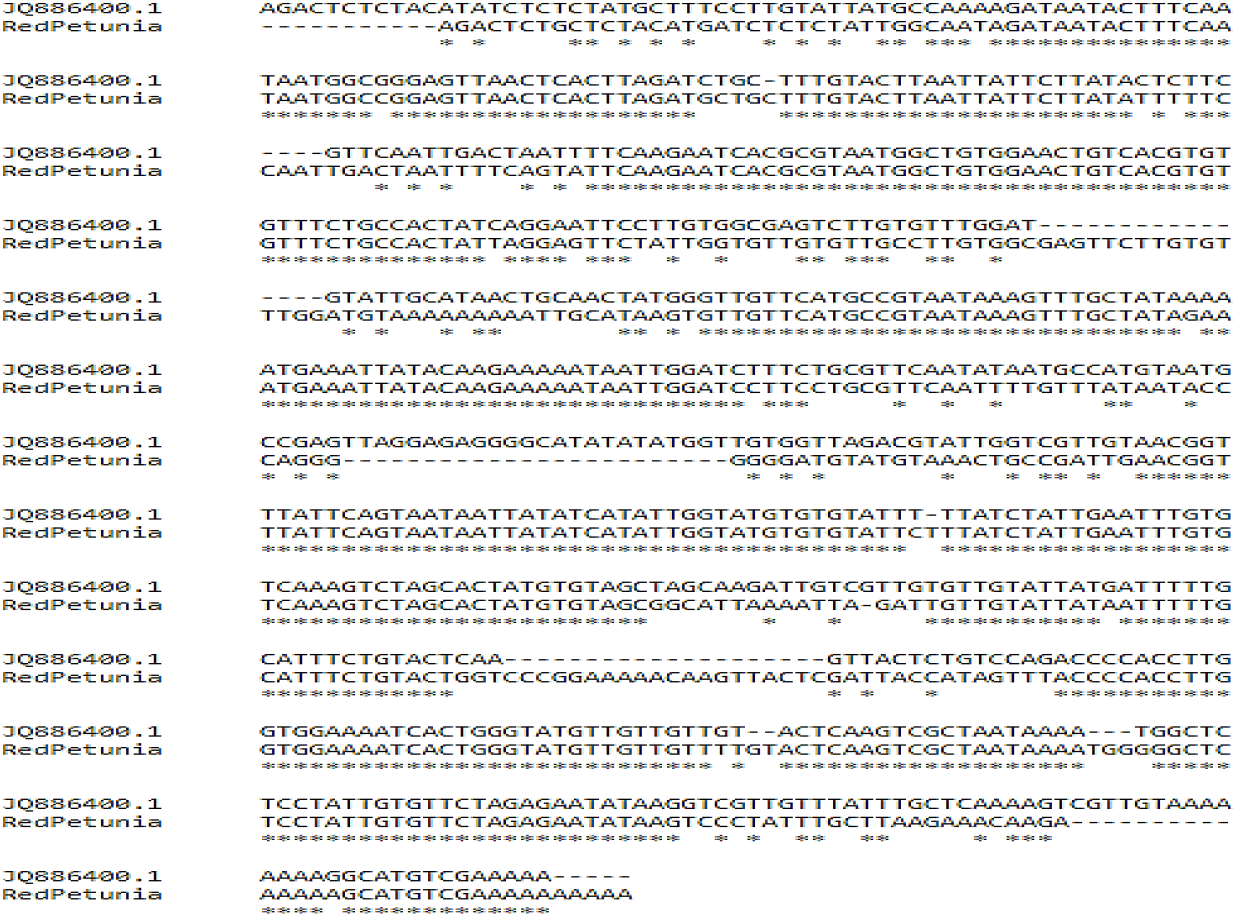
Alignment of Red *Petunia*

### Alignment of White Petunia Isolates with Reference Sequence (JQ886400)

Alignment of isolated sequence from Red Petunia with reference sequence was done and it showed 95% identity with cds of reference sequence as shown in (figure 8).

**Figure 8.**
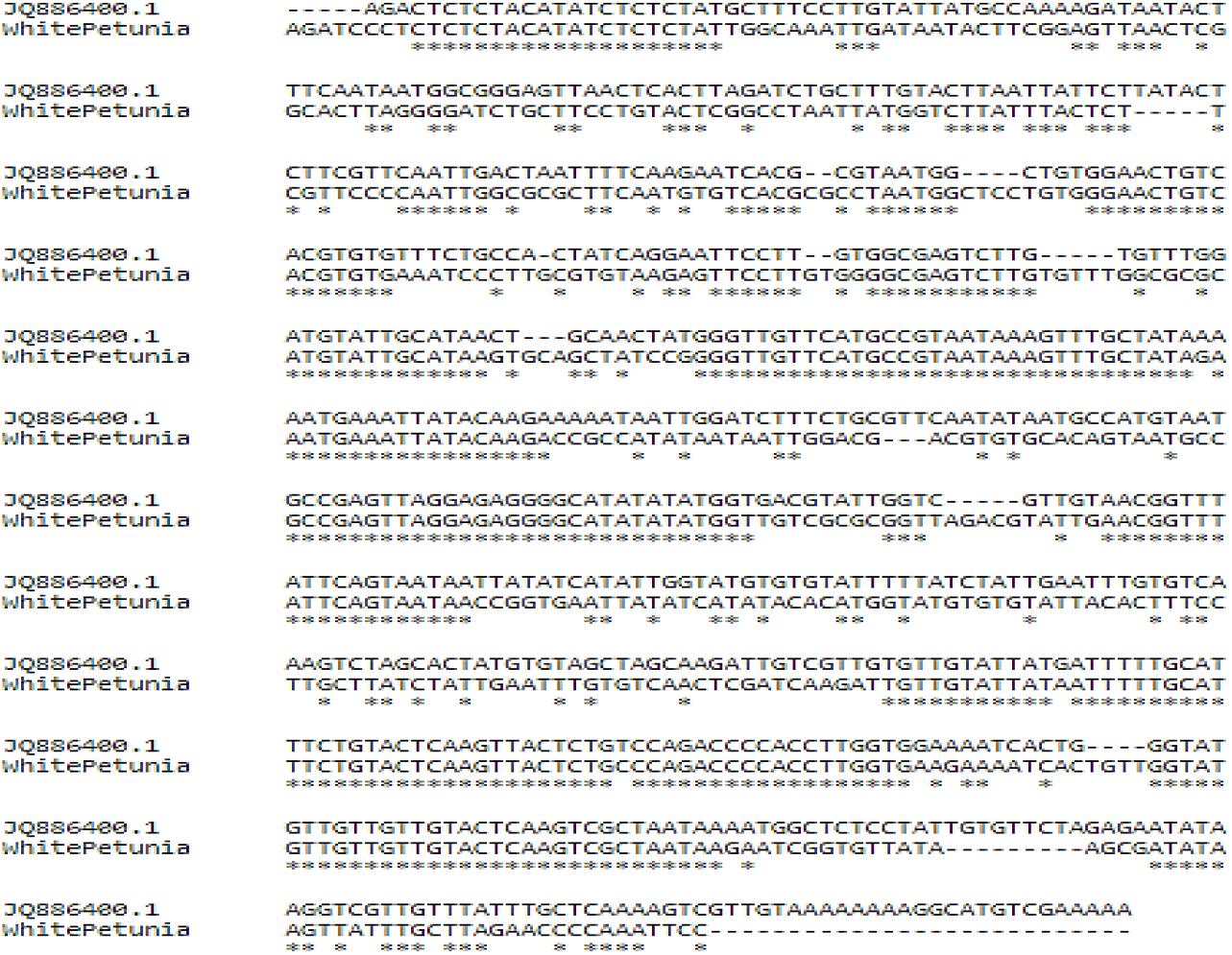
Alignment of White *Petunia*

### Representation of protein sequence of White, Blue and Red Petunia Isolates

Protein sequence of all the three selected petunia plants has been illustrated in the (figure 9) with respect to reference protein sequence AFM52764.1 which depicts its name as cyclotide precursor 3, location at 47..286, 79 amino acids, CDS position 181, protein position as 61, having protein sequence (PCGESCVWMYCITAT[M]GCSCRNKVCYKNEI). Hence, whole details about reference protein sequence are detailed in it.

**Figure 9.**
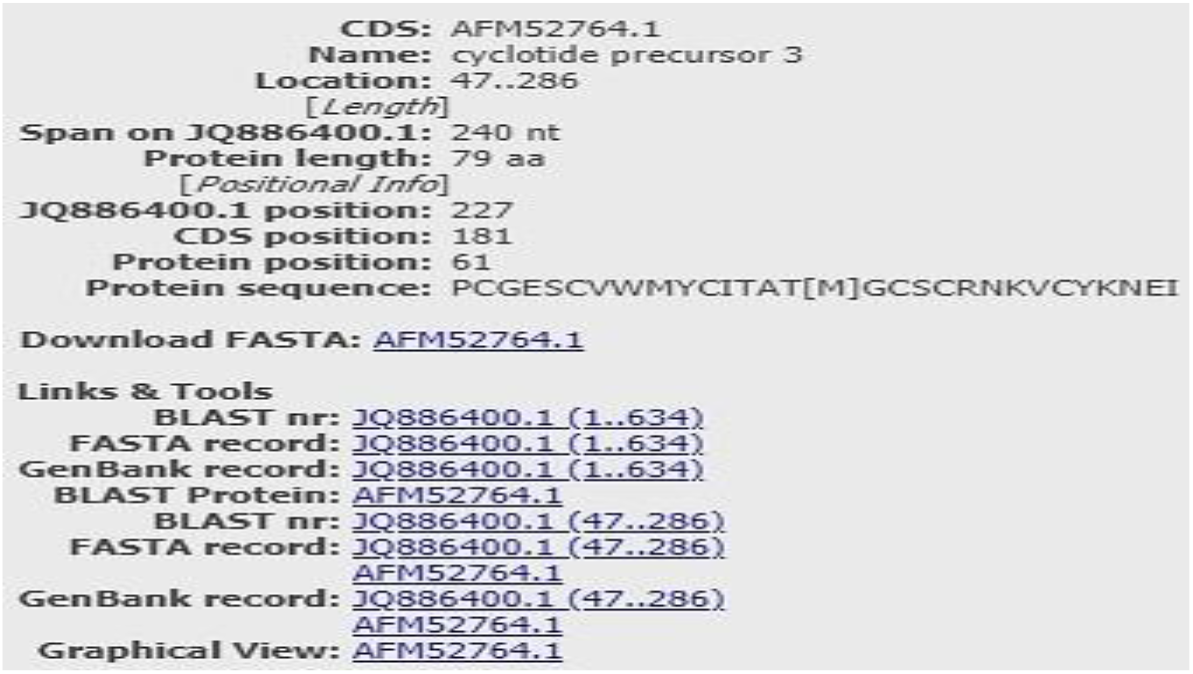
Detail of Reference Protein Sequence (AFM52764.1)

### Multiple Alignment

Multiple sequence alignment Clustal 0(1.2.4) is present in the (figure 10) which shows reference cyclotides and cyclotides of white, blue, and red petunia plants at various positions.

**Figure 10.**
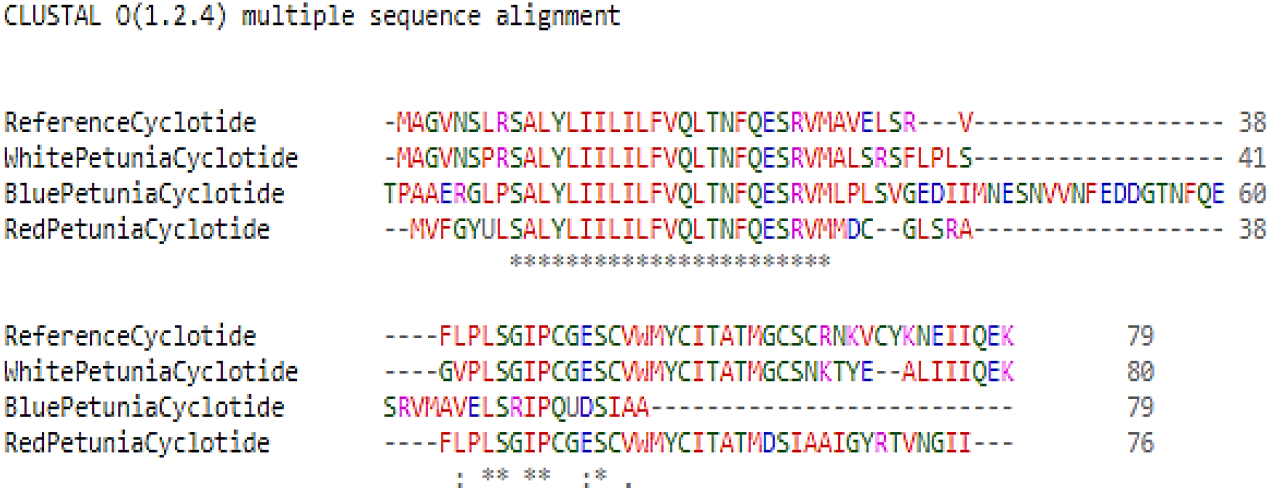
Multiple alignment of isolates

## Discussion

Cyclotides are circular plant proteins with nearly 30 amino acids that are the largest and best-studied class of cyclic peptides generated from plants and between the C-terminal and the N-terminal, an amino acid connection forms the circular peptide backbone. Six cysteine residues make up cyclotides, where each of the two cysteines forms a disulphide bond (19). Thus, the cyclic cysteine knot known as the CCK motif is generated by joining three disulphide bonds to form a cyclic structure. Cyclotides are the most stable globular microproteins because of the following disulphide linkages that form a cyclic cystine knot across the head-to-tail cyclized backbone. It is alleged that cyclotides function as protective chemicals inside plants. Thus, the application of insecticidal qualities has begun (20).

According to this research, Cyclotides are linked to the vascular attributes of petunia leaf tissues, which is compatible with previously discovered small molecules and peptidic mediators of plant resilience (4). Defensins in capsicum, trypsin inhibitors in pumpkin fruit, cysteine proteinase inhibitors in maize, glucosinolates, precursors of poisonous cyano chemicals in Arabidopsis, and terpenes implicated in squirt-gun defenses in Bursera are a few examples. Therefore, the presence of elevated cyclotide concentrations in these regions may influence herbivore feeding habits and aid in plant defense. This study expands the list of plant groups known to produce cyclotides and encourages more research into Solanaceae species for cyclotides. Based on the currently documented variance in cyclotide gene architectures, it appears probable that additional notable differences will be found in plant families that include cyclotides but have not yet been identified.

In order to comprehend its origins in evolution as the consequence of either convergent evolution or even transposable element activity, this combined information will be essential. A 1.5% agarose gel was loaded with DNA extracted from three Solanaceae plants and a PCR sample of red, white, and blue petunias to determine whether the necessary gene was present. The bands were found around roughly 690, 670, and 750 bp. The outcome is displayed in figures in results section. Furthermore, used BLAST, NCBI, and other bioinformatics tools to compare the isolated gene’s sequence with the reference sequence, once it has been received and together with the coding regions comprising three cyclotide-encoding genes (White, Blue, and Red) and the processed amino acid sequences retrieved through BLAST Petunia Cyclotide that encode precursor proteins with 79 residues are also displayed. Moreover, 90% of the DNA sequence of the Petunia x hybrid cyclotide precursor 3 mRNA is identical to that of Blue Petunia using the reference sequence likewise, Red and White Petunias exhibit 93% and 95% identity. Although it is still unclear how common cyclotides may be among Solanaceae plants, BLAST searches that Petunia x Hybrida matches were the only ones found when full-length PETUNIA sequences were utilized to query all GenBankTM nucleotide sequences, including NCBI databases (21).

Given how essential the Solanaceae plant family is to human nutrition (22), the identification of cyclic compounds in this class is noteworthy and fascinating. Additionally, unlike other drug design features, cyclotides are not impacted by biochemical, thermal, or protease processing constraints (23), because it lacks ends and specialized locations for protease attack, the circular backbone can, to some extent, offer resistance to proteases (24). Further, the entropy defect of cyclic peptide binding to the receptor is less than that of extended linear peptide binding because of heat stability (25) and epoxides that naturally exist own an extensive array of uses. Hence, numerous biological processes, such as hemolysis, hypotension, aggression, cytotoxicity, intrauterine contraction, antibacterial agents, insecticides, and anti-fouling, are caused by these compounds.

## Conclusion

In line with previously identified small molecules and peptidic mediators of plant defense, this data demonstrates that cyclotides exhibit a substantial association with the vascular parameters of petunia leaf tissues. Furthermore, this discovery contributes to the established pool of cyclotide-producing botanical groups and provides an incentive for the subsequent analysis of Solanaceae genera for cyclotides. Given the variety in cyclotide gene structures that have been described thus far, it appears probable that additional notable variations will be found in plant families that have not yet been described but contain cyclotides. Knowing their respective positions chronological origins as either a consequence of convergent evolution or possibly the activity of transposable elements will be made possible by this combined content.

## Acknowledgements

We are extremely thankful to the Faisalabad Nursery for supplying the plants necessary to conduct our investigation. Our sincere appreciation is extended to our mentors and colleagues for their excellent advice, thought-provoking conversations, and steadfast support during this project. We appreciate the bioinformatics team’s contributions because of their proficiency in interpreting and analyzing data. Last but not least, we thank the earlier researchers whose work on cyclotide genes in the Solanaceae family served as the foundation for this inquiry.

