## supplemental table 1 for "Isolation of Cyclotide Gene from Solanaceae Family and Its Bioinformatics Analysis"

**Table 1:** Primers Details

| Oligo name | Sequence (5'-3') | length | MW | Tm | GC% | nmols | ug | $A_{260}$ units |
|---|---|---|---|---|---|---|---|---|
| PET1F | GTCACGTGTGTTTCTGCCAC | 20 | 6,075 | 59.97 | 55 | 41.6 | 302.2 | 9.18 |
| PET1R | TTTTCCACCAAGGTGGGGTC | 20 | 6,119 | 60.11 | 55 | 44.3 | 325.6 | 10.85 |

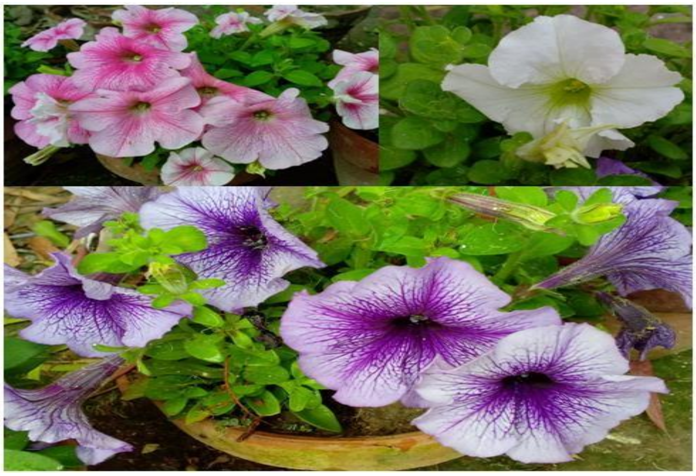

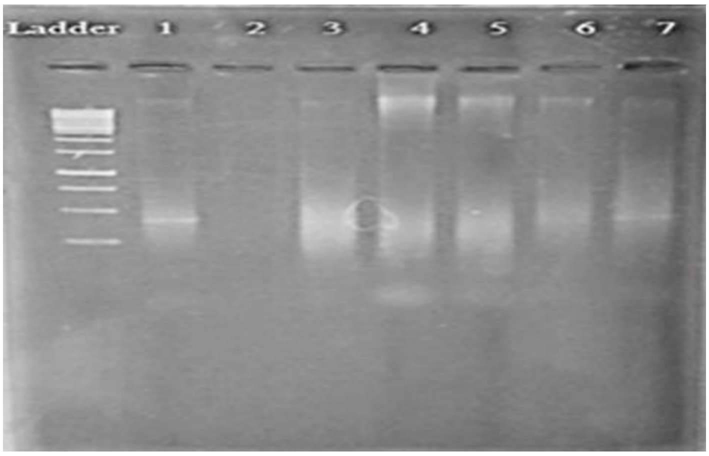

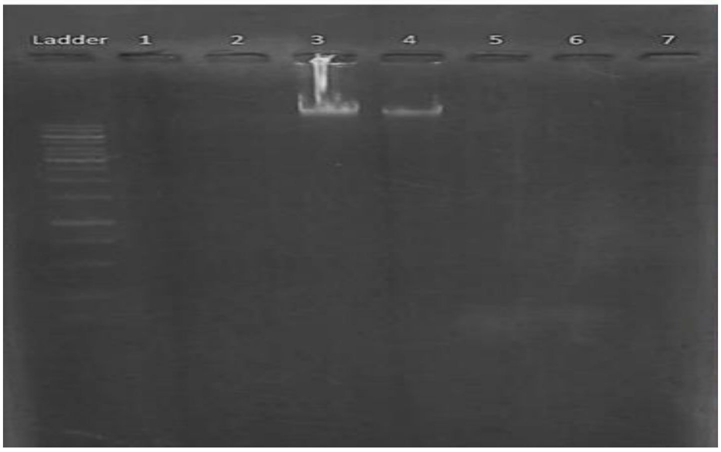


>JQ886400.1 Petunia x hybrida cyclotide precursor 3 (PETUNITIDE3) mRNA,
AGACTCTCTACATATCTCTCTATTGGCAAAGATAATACTTTCAATAATGGCGGGAGTTAACTCACTTAGA
TCTGCTTTGTACTTAATTATTCTTATACTCTTCGTTCAATTGACTAATTTTCAAGAATCACGCGTAATGG
CTGTGGAACTGTCACGTGTGTTTCTGCCACTATCAGGAATTCCTTGTGGCGAGTCTTGTGTTTGGATGTA
TTGCATAACTGCAACTATGGGTTGTTCATGCCGTAATAAAGTTTGCTATAAAAATGAAATTATACAAGAA
AAATAATTGGATCTTTCTGCGTTCAATATAATGCCATGTAATGCCGAGTTAGGAGAGGGGCATATATATG
GTTGTGGTTAGACGTATTGAACGGTTTATTCAGTAATAATTATATCATATTGGTATGTGTGTATTTTTAT
CTATTGAATTTGTGTCAAAGTCTAGCACTATGTGTAGCTAGCAAGATTGTTGTATTATGATTTTTGCATT
TCTGTACTCAAGTTACTCTGTCCAGACCCCACCTTGGTGGAAAATCACTGGGTATGTTGTTGTTGTACTC
AAGTCGCTAATAAAATGGCTCTCCTATTGTGTTCTAGAGAATATAAGTTATTTGCTCAAAAAAAAAAAAA
AAAA

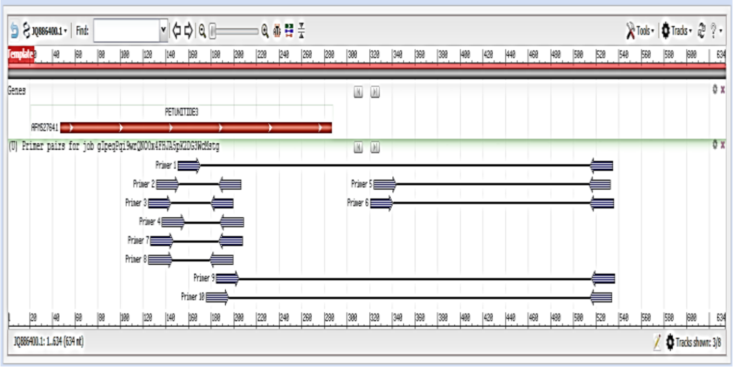


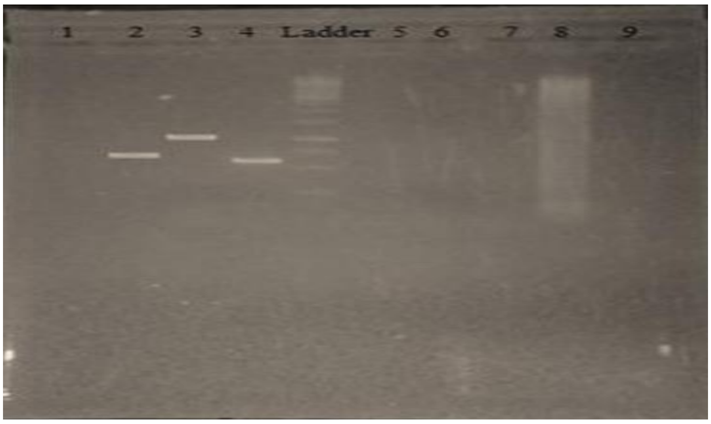

```
JQ886400.1      AGACTCTCTACATATCTCTCTATGCTTTCCTTGTATTATGCCAAAAGATAATACTTTCAA
BluePetunia     -----ATCTAGACTCTGCTC-------TGTTGATGGTGTTTCTGGGAATTGGTATT----

JQ886400.1      TAATGGCGGGAGTTAACTCACTTAGATCTGCTTTGTACTTAATTATTCTTATACTCTTCG
BluePetunia     -GCAAGTTTGATACATTCACTCCGGATCTGCTTTGTACTTAATTATTGCAGCAGAACGTG

JQ886400.1      TTCAA---------------------TTGACTAATTTTCAAGAA----------------
BluePetunia     GACTACCTGTGACAAGAGTCATCAGCCGCATGACTTTGCAACAGATCCTTGCACGTGCTG

JQ886400.1      ---------------TCACGCGTAATGGCTGTGGAACTGTCACGTGTGTTTCTGCC---A
BluePetunia     TTGGAGAAGATATAATTATGAATGAAAGTAATGAAACAGTACTTTGGTTAACTTTGAGGA

JQ886400.1      CTATCAGGAATTCCTTGTGGCGAGTCTTGTGTTTGGATGTATTGCATAACTGCAACTATG
BluePetunia     TGATGGGGACAAGGTTAGTGTGACTCTTGAAAATGGACAGCAATATACTGGTGATCTTCT

JQ886400.1      GGTTGTTCATGCCGTAATAAAGTTTGCTATAAAAATGAAATTATACAAGAAAAATAATTG
BluePetunia     GGTTGGTGCTGATGGCATACGGTCTAAGGTAAGGACAAATTTGTTCGGACCAAGC-----

JQ886400.1      GATCTTTCTGCGTTCAATATAATGCCATGTAATGCCGAGTTAGGAGAGGG-----GCATA
BluePetunia     -GAAGCTAATTATTCAATATAATGCCATGTGGCTACACATGTTACACTGGAATTTGCAAT

JQ886400.1      TATATGGTTGTGGTTAGACGTATTGGTCGTTGTAACGGTTT-------------------
BluePetunia     TTTATTCCTGCTGATATTGAGACAGTTGGGTACCGCGTCTTTTTGGGGCACAAACAGTAC

JQ886400.1      -------------------ATTCAGTAATAATTATATCATATTGGTATGTGTGTATTTTT
BluePetunia     TTTGTTTCTTCGGATGTTGGTGCTGGAAAGATGCAGTAGGAGAGGGGCATATATATGGTT

JQ886400.1      ATCTATTGAATTTGTGTCAAAGTCTAGC---ACTATGTGTAGCT------AGCAAGATTG
BluePetunia     GT----GGTATGCATTTCATAATGAACCAGCTGGCGGTGTGGATGGTCCAAACGGTATGA

JQ886400.1      TCGTTGTGTTGTATTATGATTTTTGCA-TTTCTGTACTCAAGTTACTCTGTCCAGACCCC
BluePetunia     AGGCAAGATTGCTTGAAATATTTGGCGGTTGGTGTGACAATGTTATAGACCTATTAGTTG

JQ886400.1      ACCTTGGTGGAAAATCACTGGGTATGTTGTTGTTGTACTCAAGTCGCTAATAAAATGGCT
BluePetunia     CCACTGATGAAGAAGCAATTC----------TTCGACGTGACATCTACGA-----TAGAG

JQ886400.1      CTCCTATTGTGTTCTAGAGAATATAAGGTCGTTGTTTATTTG-CTCAAAAGTCGTTGTAA
BluePetunia     CGCCAACCTTTAGTTGGGGAAGAGGCCGTGTTACATTGCTTGGGGACTCTGTCCATGCTA

JQ886400.1      AAAAAAGGCATGTCGAAAAA--------------
BluePetunia     TGCAGCCTAATTTGGGACAAGGTGGTTGCATGGC
```

```
JQ886400.1      AGACTCTCTACATATCTCTCTATGCTTTCCTTGTATTATGCCAAAAGATAATACTTTCAA
RedPetunia      -----------AGACTCTGCTCTACATGATCTCTCTATTGGCAATAGATAATACTTTCAA

JQ886400.1      TAATGGCGGGAGTTAACTCACTTAGATCTGC-TTTGTACTTAATTATTCTTATACTCTTC
RedPetunia      TAATGGCCGGAGTTAACTCACTTAGATGCTGCTTTGTACTTAATTATTCTTATATTTTTC

JQ886400.1      ----GTTCAATTGACTAATTTTCAAGAATCACGCGTAATGGCTGTGGAACTGTCACGTGT
RedPetunia      CAATTGACTAATTTTCAGTATTCAAGAATCACGCGTAATGGCTGTGGAACTGTCACGTGT

JQ886400.1      GTTTCTGCCACTATCAGGAATTCCTTGTGGCGAGTCTTGTGTTTGGAT------------
RedPetunia      GTTTCTGCCACTATTAGGAGTTCTATTGGTGTTGTGTTGCCTTGTGGCGAGTTCTTGTGT

JQ886400.1      ----GTATTGCATAACTGCAACTATGGGTTGTTCATGCCGTAATAAAGTTTGCTATAAAA
RedPetunia      TTGGATGTAAAAAAAAATTGCATAAGTGTTGTTCATGCCGTAATAAAGTTTGCTATAGAA

JQ886400.1      ATGAAATTATACAAGAAAAATAATTGGATCTTTCTGCGTTCAATATAATGCCATGTAATG
RedPetunia      ATGAAATTATACAAGAAAAATAATTGGATCCTTCCTGCGTTCAATTTTGTTTATAATACC

JQ886400.1      CCGAGTTAGGAGAGGGGCATATATATGGTTGTGGTTAGACGTATTGGTCGTTGTAACGGT
RedPetunia      CAGGG------------------------GGGGATGTATGTAAACTGCCGATTGAACGGT

JQ886400.1      TTATTCAGTAATAATTATATCATATTGGTATGTGTGTATTT-TTATCTATTGAATTTGTG
RedPetunia      TTATTCAGTAATAATTATATCATATTGGTATGTGTGTATTCTTTATCTATTGAATTTGTG

JQ886400.1      TCAAAGTCTAGCACTATGTGTAGCTAGCAAGATTGTCGTTGTGTTGTATTATGATTTTTG
RedPetunia      TCAAAGTCTAGCACTATGTGTAGCGGCATTAAAATTA-GATTGTTGTATTATAATTTTTG

JQ886400.1      CATTTCTGTACTCAA--------------------GTTACTCTGTCCAGACCCCACCTTG
RedPetunia      CATTTCTGTACTGGTCCCGGAAAAACAAGTTACTCGATTACCATAGTTTACCCCACCTTG

JQ886400.1      GTGGAAAATCACTGGGTATGTTGTTGTTGT--ACTCAAGTCGCTAATAAAA---TGGCTC
RedPetunia      GTGGAAAATCACTGGGTATGTTGTTGTTTTGTACTCAAGTCGCTAATAAAATGGGGGCTC

JQ886400.1      TCCTATTGTGTTCTAGAGAATATAAGGTCGTTGTTTATTTGCTCAAAAGTCGTTGTAAAA
RedPetunia      TCCTATTGTGTTCTAGAGAATATAAGTCCCTATTTGCTTAAGAAACAAGA----------

JQ886400.1      AAAAGGCATGTCGAAAAA-----
RedPetunia      AAAAAGCATGTCGAAAAAAAAAA
```

```
JQ886400.1         -----AGACTCTCTACATATCTCTCTATGCTTTCCTTGTATTATGCCAAAAGATAATACT
WhitePetunia       AGATCCCTCTCTCTACATATCTCTCTATTGGCAAATTGATAATACTTCGGAGTTAACTCG
                           ********************       ***            ** ***  *

JQ886400.1         TTCAATAATGGCGGGAGTTAACTCACTTAGATCTGCTTTGTACTTAATTATTCTTATACT
WhitePetunia       GCACTTAGGGGATCTGCTTCCTGTACTCGGCCTAATTATGGTCTTATTTACTCT-----T
                        **  **      **     ***  *      * **  **** *** ***     *

JQ886400.1         CTTCGTTCAATTGACTAATTTTCAAGAATCACG--CGTAATGG----CTGTGGAACTGTC
WhitePetunia       CGTTCCCCAATTGGCGCGCTTCAATGTGTCACGCGCCTAATGGCTCCTGTGGGAACTGTC
                   * *    ****** *    **  * *  *****  * ******        *********

JQ886400.1         ACGTGTGTTTCTGCCA-CTATCAGGAATTCCTT--GTGGCGAGTCTTG-----TGTTTGG
WhitePetunia       ACGTGTGAAATCCCTTGCGTGTAAGAGTTCCTTGTGGGGCGAGTCTTGTGTTTGGCGCGC
                   *******      *   *    * ** ******  * ***********      *   *

JQ886400.1         ATGTATTGCATAACT---GCAACTATGGGTTGTTCATGCCGTAATAAAGTTTGCTATAAA
WhitePetunia       ATGTATTGCATAAGTGCAGCTATCCGGGGTTGTTCATGCCGTAATAAAGTTTGCTATAGA
                   ************* *   ** *    ******************************** *

JQ886400.1         AATGAAATTATACAAGAAAAATAATTGGATCTTTCTGCGTTCAATATAATGCCATGTAAT
WhitePetunia       AATGAAATTATACAAGACCGCCATATAATAATTGGACG---ACGTGTGCACAGTAATGCC
                   *****************     *  *     **           * *         *

JQ886400.1         GCCGAGTTAGGAGAGGGGCATATATATGGTGACGTATTGGTC-----GTTGTAACGGTTT
WhitePetunia       GCCGAGTTAGGAGAGGGGCATATATATGGTTGTCGCGCGGTTAGACGTATTGAACGGTTT
                   ******************************        ***        *  ********

JQ886400.1         ATTCAGTAATAATTATATCATATTGGTATGTGTGTATTTTTATCTATTGAATTTGTGTCA
WhitePetunia       ATTCAGTAATAACCGGTGAATTATATCATATACACATGGTATGTGTGTATTACACTTTCC
                   ************       **  *   ** *    **  *       *       * **

JQ886400.1         AAGTCTAGCACTATGTGTAGCTAGCAAGATTGTCGTTGTGTTGTATTATGATTTTTGCAT
WhitePetunia       TTGCTTATCTATTGAATTTGTGTCAACTCGATCAAGATTGTTGTATTATAATTTTTGCAT
                     *  ** *  *     * *     *            *********** **********

JQ886400.1         TTCTGTACTCAAGTTACTCTGTCCAGACCCCACCTTGGTGGAAAATCACTG----GGTAT
WhitePetunia       TTCTGTACTCAAGTTACTCTGCCCAGACCCCACCTTGGTGAAGAAAATCACTGTTGGTAT
                   ********************* ****************** * **   *      *****

JQ886400.1         GTTGTTGTTGTACTCAAGTCGCTAATAAAATGGCTCTCCTATTGTGTTCTAGAGAATATA
WhitePetunia       GTTGTTGTTGTACTCAAGTCGCTAATAAGAATCGGTGTTATA---------AGCGATATA
                   **************************** *                         *****

JQ886400.1         AGGTCGTTGTTTATTTGCTCAAAAGTCGTTGTAAAAAAAAGGCATGTCGAAAAA
WhitePetunia       AGTTATTTGCTTAGAACCCCAAATTCC---------------------------
                   ** *  *** ***    * ****   *
```

**CDS:** AFM52764.1
**Name:** cyclotide precursor 3
**Location:** 47..286
[*Length*]
**Span on JQ886400.1:** 240 nt
**Protein length:** 79 aa
[*Positional Info*]
**JQ886400.1 position:** 227
**CDS position:** 181
**Protein position:** 61
**Protein sequence:** PCGESCVWMYCITAT[M]GCSCRNKVCYKNEI

**Download FASTA:** AFM52764.1

**Links & Tools**
**BLAST nr:** JQ886400.1 (1..634)
**FASTA record:** JQ886400.1 (1..634)
**GenBank record:** JQ886400.1 (1..634)
**BLAST Protein:** AFM52764.1
**BLAST nr:** JQ886400.1 (47..286)
**FASTA record:** JQ886400.1 (47..286)
AFM52764.1
**GenBank record:** JQ886400.1 (47..286)
AFM52764.1
**Graphical View:** AFM52764.1

```
CLUSTAL O(1.2.4) multiple sequence alignment


ReferenceCyclotide          -MAGVNSLRSALYLIILILFVQLTNFQESRVMAVELSR---V------------------ 38
WhitePetuniaCyclotide       -MAGVNSPRSALYLIILILFVQLTNFQESRVMALSRSFLPLS------------------ 41
BluePetuniaCyclotide        TPAAERGLPSALYLIILILFVQLTNFQESRVMLPLSVGEDIIMNESNVVNFEDDGTNFQE 60
RedPetuniaCyclotide         --MVFGYULSALYLIILILFVQLTNFQESRVMMDC--GLSRA------------------ 38
                                     ***********************


ReferenceCyclotide          ----FLPLSGIPCGESCVWMYCITATMGCSCRNKVCYKNEIIQEK        79
WhitePetuniaCyclotide       ----GVPLSGIPCGESCVWMYCITATMGCSNKTYE--ALIIIQEK        80
BluePetuniaCyclotide        SRVMAVELSRIPQUDSIAA--------------------------        79
RedPetuniaCyclotide         ----FLPLSGIPCGESCVWMYCITATMDSIAAIGYRTVNGII---        76
                                 : ** **  :* .
```
